# RNA sequence design and protein–DNA specificity prediction with NA-MPNN

**DOI:** 10.1101/2025.10.03.679414

**Authors:** Andrew Kubaney, Andrew Favor, Lilian McHugh, Raktim Mitra, Robert Pecoraro, Justas Dauparas, Cameron Glasscock, David Baker

## Abstract

RNA sequence design and protein–DNA binding specificity prediction can both be framed as nucleic acid inverse-folding problems: finding the most likely nucleic acid sequences given a fixed three-dimensional structure of a nucleic acid or nucleic acid–protein complex. While task-specific tools have been developed, no unified deep learning model for nucleic acid inverse folding has been described; a single model would have larger and more diverse datasets available for training and a considerably greater range of applicability. Here we introduce Nucleic Acid MPNN (NA-MPNN), a message-passing neural network that treats proteins, DNA, and RNA within a unified biopolymer graph representation. NA-MPNN outperforms previous methods on RNA sequence design and fixed-dock protein–DNA specificity prediction, and should be broadly useful for *de novo* RNA structure design and prediction of DNA-binding specificity.

## Introduction

The design of nucleic acid sequences with specific structural or functional properties has considerable potential for the creation of aptamers^1^, ribozymes^2^, CRISPR guide RNA scaffolds^3,4^, and non-native ribosomes^5^. Likewise, accurate prediction of protein–DNA binding specificity is essential for understanding and manipulating processes such as transcriptional regulation^6–11^, DNA repair^12^, and genome editing^13,14^. Together, these two challenges—RNA sequence design and protein–DNA specificity prediction—define complementary facets of nucleic acid inverse folding: the latter in the context of a fixed protein structure and the former without external context. Current methods for nucleic acid inverse folding are task-specific and leave room for accuracy improvement. For RNA, backbone-conditioned inverse folding has been addressed by geometric deep-learning approaches such as gRNAde^15^ and RhoDesign^16^. Protein–DNA specificity can be estimated either in a “dock-free” setting—employing structure-prediction engines such as RoseTTAFoldNA (RFNA)^17^, RoseTTAFold All-Atom (RFAA)^18^, AlphaFold 3^19^, MELD-DNA^20^, or family-specific models including rCLAMPS^11^—or in a “fixed-dock” setting, where the complex geometry is given and methods like DeepPBS^21^ infer base preferences directly. For training accurate deep learning models, the availability of large and diverse training sets is critical. For the RNA sequence-design problem, the limited availability of high-resolution RNA structures presents a significant hurdle: as of January 2025, the Protein Data Bank (PDB)^22,23^ includes 8,961 RNA-containing entries compared to 235,538 protein-containing entries, a disparity that complicates model training and benchmarking. For protein–DNA binding specificity prediction, in addition to co-crystal structures, assays such as SELEX or PBM have yielded per-position base specificity preferences^24–27^. Despite the fact that both RNA sequence design and fixed-dock protein–DNA specificity prediction can be framed as nucleic acid inverse-folding problems, no single neural architecture has yet been developed to address both tasks simultaneously.

We hypothesized that a unified deep-learning–based nucleic acid inverse-folding model could address both problems, with potential advantages from jointly training on the disparate datasets available in the two cases, and a considerably broader range of applicability. Inspired by the accuracy and widespread usage of the deep learning ProteinMPNN^28^ network and its generalization to ligand-aware contexts in LigandMPNN^29^, we set out to develop a message-passing neural network capable of processing both nucleic acid and protein backbones.

## Results

### NA-MPNN Architecture

We reasoned that we could develop a general biopolymer sequence-design (inverse-folding) network by extending ProteinMPNN to enable computation over nucleic acid backbones (Fig. 1a). ProteinMPNN has a graph neural-network architecture with residues as nodes; in both ProteinMPNN and LigandMPNN, the nodes can only be protein residues. We created a new model, called NA-MPNN, in which the nodes can be either protein residues or DNA or RNA bases, and edges between nodes can be protein–protein, nucleic acid–nucleic acid, or protein–nucleic acid. Edges are determined via the Cα atom of proteins and the C1′ atom of nucleic acids. Following ProteinMPNN, we construct a virtual Cβ for all protein residues; for nucleic acids we add an analogous virtual side-chain N atom. The edge features are radial-basis-function embeddings of all pairwise backbone atom distances between the two residues or bases; for proteins the N, Cα, C, O, and virtual Cβ atoms, and for nucleic acids, the phosphate atoms (P, OP1, OP2), sugar atoms (C1′–5′, O3′–5′, O2′ for RNA), and virtual side-chain N atom (Fig. 1b). Each node exchanges messages with its 32 nearest neighbors (ProteinMPNN uses 48), and small isotropic Gaussian noise (σ = 0.1 Å) is injected during training to discourage overfitting to crystal artifacts. As in ProteinMPNN, label smoothing across residue tokens is applied to reduce model overconfidence^30^, but the probability mass is redistributed within polymer classes to prevent leakage across protein and nucleic acid alphabets. The random-order autoregressive decoder enables partial sequence fixation during design (Fig. 2a). The encoder and decoder architectures mirror those of ProteinMPNN and LigandMPNN (Fig. 1c).

**Figure 1.**
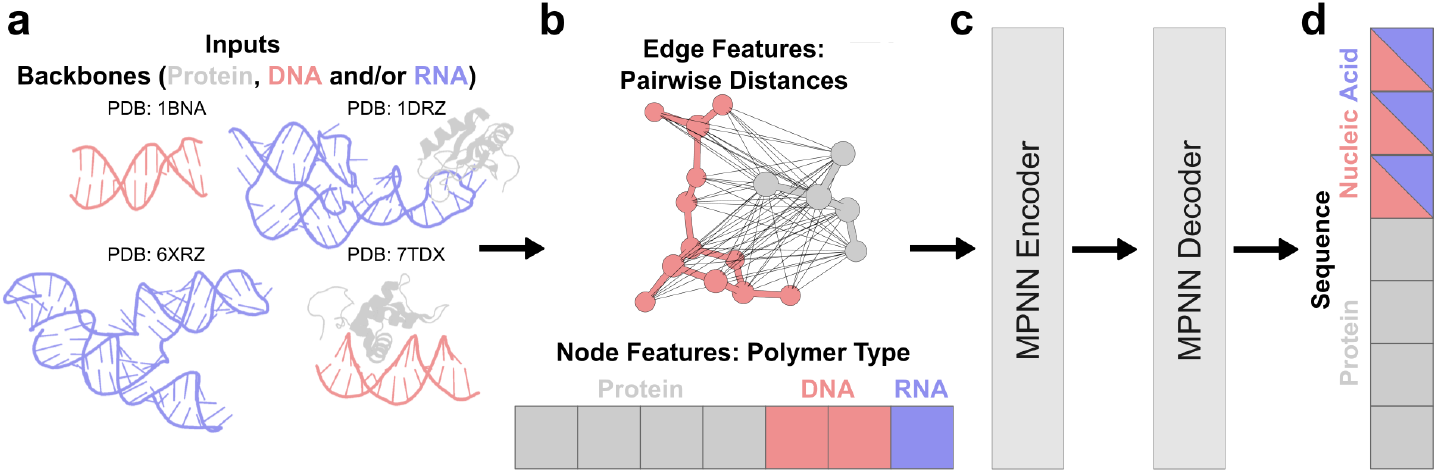
NA-MPNN architecture. The schematic illustrates the message-passing neural network that underpins NA-MPNN. **(a)** Protein, DNA, and/or RNA backbones are supplied as input graphs. **(b)** Nodes correspond to residues annotated with a one-hot polymer-type embedding (protein, DNA, RNA), and edges connect the 32 nearest neighbors of each residue, and their features comprise radial-basis-function embeddings of all inter-residue backbone distances, including a virtual first side-chain atom for both proteins and nucleic acids. **(c)** The resulting node- and edge-feature tensors are processed by the standard MPNN encoder–decoder stack of ProteinMPNN. **(d)** The network outputs categorical distributions over 21 amino-acid tokens and five shared nucleic acid tokens (DA/A, DC/C, DG/G, DT/U, DX/RX).

**Figure 2.**
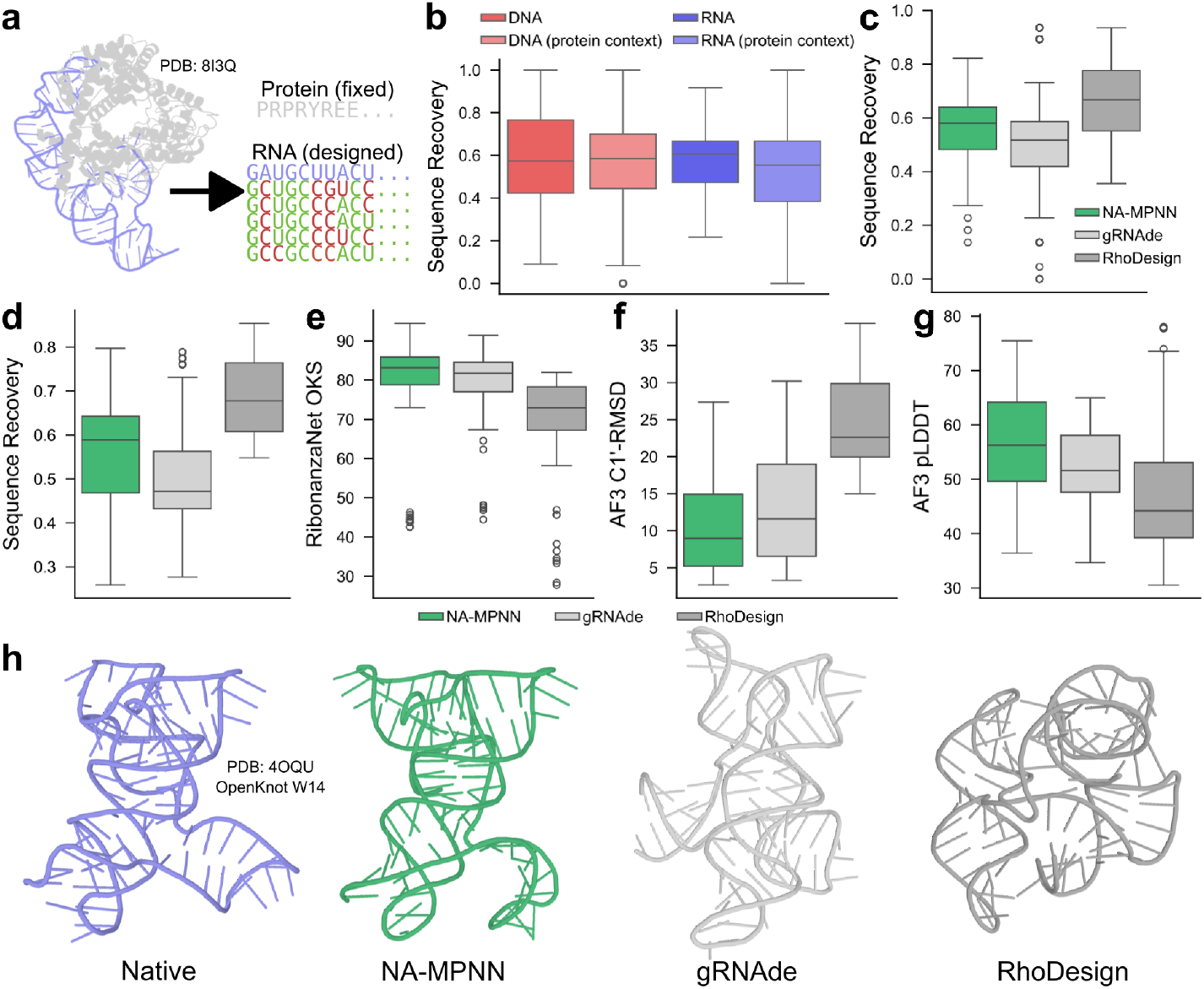
*In silico* sequence-design performance of NA-MPNN. **(a)** An example of protein-conditioned RNA inverse folding: the protein sequence is held fixed while NA-MPNN designs the RNA chain. Designed sequences (rows) are aligned beneath the native sequence (top row); matches are colored green, mismatches red. **(b)** Distribution of sequence recovery on the design test set, broken down by four contexts—DNA-only, RNA-only, DNA in protein context, and RNA in protein-context—showing comparable performance across classes with the highest median for standalone RNA. **(c)** Sequence recovery comparison on the RNA-monomer test set: NA-MPNN surpasses gRNAde but is outperformed by RhoDesign. **(d)** Distribution of sequence recovery on the pseudoknot test set, demonstrating a similar trend to the RNA-monomer test set. **(e)** OpenKnot score (OKS) of the RibonanzaNet-predicted reactivity profiles for the designed pseudoknot sequences, demonstrating that NA-MPNN achieves superior secondary structure recovery. **(f–g)** C1′-RMSD and pLDDT distributions derived from AlphaFold 3 predictions of designed pseudoknot sequences, highlighting that NA-MPNN sequences are more likely to confidently adopt the desired tertiary structure. **(h)** Qualitative comparison of AlphaFold 3-predicted tertiary structure for one pseudoknot target: the native structure and the best (lowest C1′-RMSD) of ten NA-MPNN, gRNAde, and RhoDesign designs are shown; NA-MPNN most closely recapitulates the target.

NA-MPNN departs from ProteinMPNN in two architectural respects. First, instead of zero initial node features, each residue receives an explicit one-hot polymer-type embedding (protein, DNA, RNA) (Fig. 1b); although the network could in principle infer polymer identity from the backbone atoms that are present, providing this directly accelerates learning. Second, the token alphabet is expanded to cover nucleic acids in a unified manner: a single token is shared between the deoxyribo- and ribo-forms of each canonical base (DA/A, DC/C, DG/G, DT/U) as well as an unknown nucleic acid token (DX/RX) (Fig. 1d). This collapsed alphabet encourages cross-learning between DNA and RNA contexts—particularly valuable because NA-MPNN does not see side-chain chemistry—and in our experiments improved sequence recovery on held-out validation sets.

Although NA-MPNN uses a unified graph architecture for all polymers (protein, DNA, RNA), we train two task-specific models: a design model for backbone-conditioned sequence design and a specificity model for fixed-dock protein–DNA binding preferences. Both models are optimized with per-position cross-entropy, but differ in their supervision targets. For design, the target is the crystallographic sequence encoded as a label-smoothed one-hot vector at each nucleic acid position, encouraging the network to propose a single cohesive sequence that best fits the input backbone (and any protein context). For specificity, the target is an empirical position probability matrix (PPM) when available, so the cross-entropy loss enforces distribution matching between the model’s predicted base probabilities and the experimental base frequencies at each position. We adopt separate trainings because PPMs intentionally marginalize higher-order couplings between positions that may be essential for designing a single realizable sequence, whereas the specificity objective aims precisely at these marginal preferences near protein–DNA interfaces.

### Data Augmentation

The specificity model utilizes a set of data augmentations tailored to learning position probability matrix (PPM) targets. Whenever an experimental PPM is available for a DNA residue, we replace the one-hot ground-truth label in the negative log-likelihood loss with that PPM column, exploiting the fact that both tensors share the same shape and semantics. For nucleic acid-only complexes—that is, structures lacking protein context—we impose a uniform PPM (0.25 probability for each of the four canonical bases) at every position. In protein–DNA/RNA complexes we stochastically drop the protein chains with 50% probability; when the protein is removed, all nucleic acid positions are likewise assigned the uniform PPM (Fig. 4a). Finally, even in the presence of protein we overwrite positions that lie > 5 Å (heavy-atom distance) from any side-chain atom of the protein and have no experimental PPM with the uniform PPM. Collectively, these interventions helped the network to focus on sequence preferences that are induced by protein contacts rather than artifacts of nucleic acid backbone geometry, while simultaneously permitting the inclusion of examples that lack experimental PPMs by treating non-interface positions as information-free.

### Datasets

Both NA-MPNN variants were trained on PDB entries containing at least one nucleic acid chain and resolved to ≤ 3.5 Å (or with no resolution reported, e.g., NMR) and with a total polymer length ≤ 6,000 residues. The experimental PPMs used by DeepPBS^21^ were associated with the corresponding crystal structures. The specificity model was augmented with RFNA/RFAA predictions distilled from CIS-BP^26^ and TRANSFAC^27^, giving access to a further 27,546 structures with matched PPMs after filtering. For design, nucleic acid chains were clustered at 80% sequence identity, and entire nucleic acid sequence clusters were held out as a test set. A separate pseudoknot subset (10 clusters) and an RNA-monomer subset were carved from the test split. The resulting design test set comprises 39 DNA-only, 88 RNA-only, 237 protein–DNA and 139 protein–RNA complexes; the RNA-monomer and pseudoknot subsets contain 63 and 10 examples, respectively. For specificity, protein chains were clustered at 40% sequence identity, and entire protein sequence clusters were held out as a test set. The test set contained 228 structures from RFNA/RFAA distillation with experimentally determined PPMs. During sequence-design evaluation, each backbone was sampled ten times with temperature = 0.1, and all performance metrics were aggregated over the distribution of sequences and backbones. For specificity assessment, the tuned configuration of 30 samples at temperature = 0.6 was used. In all cases an example was retained only if every chain belonged to the requisite hold-out clusters, ensuring strict sequence-level independence across splits.

### Model Performance on Nucleic Acid Sequence Design

NA-MPNN achieves robust sequence recovery across both standalone nucleic acids and protein-bound contexts (Fig. 2b), with median recoveries of 57.4% for DNA-only, 60.5% for RNA-only, 58.6% for DNA in protein context, and 55.4% for RNA in protein context. On the RNA-monomer set NA-MPNN (median recovery = 58.0%) outperforms gRNAde^15^ (median = 51.7%) yet trails RhoDesign^16^ (median = 66.7%) (Fig. 2c), a trend that persists on the pseudoknot subset (median = 58.9% vs. gRNAde 47.2% and RhoDesign 67.7%) (Fig. 2d). Crucially, NA-MPNN designs exhibit superior structural fidelity: on pseudoknots the model attains higher predicted OpenKnot scores from RibonanzaNet^31^ (Fig. 2e), lower C1′-RMSD to the native structure after AlphaFold 3 prediction, and higher AlphaFold 3 pLDDT than either competitor method (median OpenKnot score = 83.2 vs. gRNAde 81.7 and RhoDesign 72.9; median C1′-RMSD = 9.0 Å vs. 11.6 Å / 22.6 Å; median pLDDT = 56.2 vs. 51.6 / 44.2) (Fig. 2f–h). These results indicate that NA-MPNN sequences are predicted to adopt the intended secondary and tertiary folds with greater accuracy and confidence than alternative backbone-conditioned RNA-design approaches.

We next assessed NA-MPNN in the community-wide OpenKnot RNA pseudoknot design challenge^31^. Round 6 of the challenge comprised 12 fixed-backbone targets; for each backbone we submitted between 10 and 15 NA-MPNN designs. Following synthesis, each design was profiled by SHAPE-Seq^31,32^; the resulting reactivity maps were converted to experimental OpenKnot scores^31^, a quantitative measure of agreement between the folded RNA and the target secondary structure (0–100, higher is better). Figure 3b–d illustrate one representative backbone together with per-design SHAPE-Seq profiles and OpenKnot scores for starting (wild-type), NA-MPNN, and gRNAde sequences. Across all 12 puzzles, NA-MPNN achieved a median experimental OpenKnot score (89.9) equivalent to or higher than gRNAde (80.7), the starting sequences (87.0), and individual Eterna player^31^ submissions (89.4) (Fig. 3a), demonstrating that the model reliably produces RNA sequences that fold into the intended secondary structure topologies.

**Figure 3.**
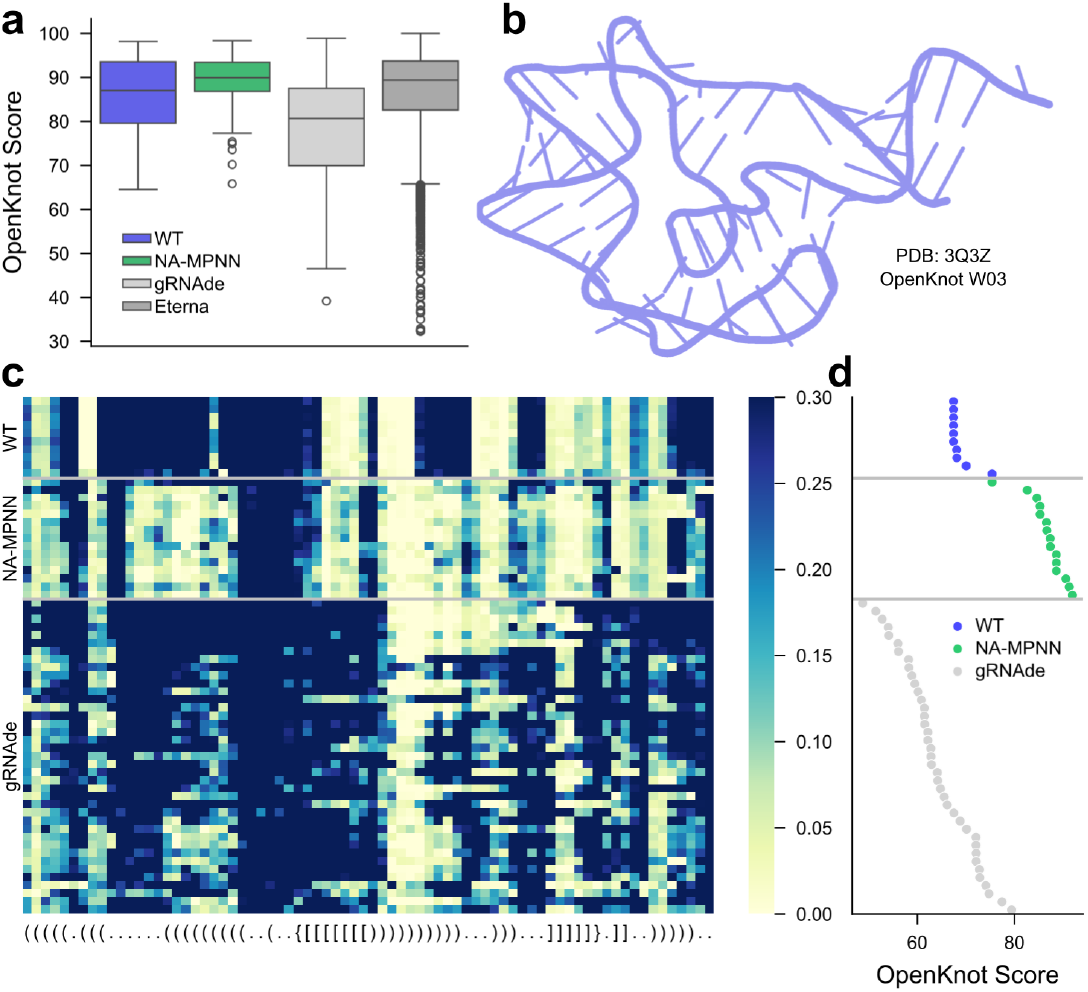
Experimental validation in the OpenKnot competition. **(a)** Distribution of experimentally derived OpenKnot scores across all puzzles in OpenKnot Round 6 for the starting sequences (WT), NA-MPNN sequences, gRNAde sequences, and Eterna player-created sequences. NA-MPNN achieves the highest median score. **(b)** Starting structure for the OpenKnot W03 puzzle. **(c)** SHAPE-Seq reactivity heat maps for the starting, NA-MPNN, and gRNAde sequences for the W03 puzzle from OpenKnot Round 6; low reactivity (yellow) denotes base-paired regions, high reactivity (blue) denotes loops. The target secondary structure is provided at the bottom. **(d)** The corresponding OpenKnot scores for the designs from the heatmap, demonstrating that NA-MPNN’s sequences achieve experimental reactivity profiles that more closely resemble the target secondary structure.

### Model Performance on Fixed-Dock Protein–DNA Specificity Prediction

For fixed-dock protein–DNA specificity, we evaluate NA-MPNN and DeepPBS^21^ on the test set of 228 protein–DNA RFNA/RFAA distillation complexes with experimental reference motifs. For NA-MPNN, we remove the DNA sequence from the input while retaining the protein sequence, protein backbone, and DNA backbone. We then perform random autoregressive decoding 30 times at temperature = 0.6, save the per-position categorical probabilities from each run, and average them to obtain a predicted PPM for every DNA position. DeepPBS predictions are generated on the same distillation complexes. Experimental PPMs are aligned to each test complex using the modified DeepPBS alignment algorithm^21^ employed during training (see Methods).

We report two per-position metrics, averaged along the aligned window for each complex: mean absolute error (MAE) and cross-entropy between the predicted and reference distributions (formulas in Methods). Conceptually, MAE measures the absolute probability error at each position; cross-entropy quantifies distributional disagreement, giving more weight to positions with higher ground-truth information content. Illustrative examples are shown in Fig. 4b, where NA-MPNN better recapitulates experimentally validated base preferences and information content than DeepPBS. Over the test set, NA-MPNN attains lower median MAE (0.52 vs. 0.87 on CIS-BP; 0.53 vs. 0.85 on TRANSFAC; 0.53 vs. 0.86 overall) and lower median cross-entropy (0.97 vs. 1.45 on CIS-BP; 1.05 vs. 1.43 on TRANSFAC; 1.00 vs. 1.44 overall) than DeepPBS (Fig. 4c–d), establishing a new state of the art for fixed-dock protein–DNA specificity prediction.

**Figure 4.**
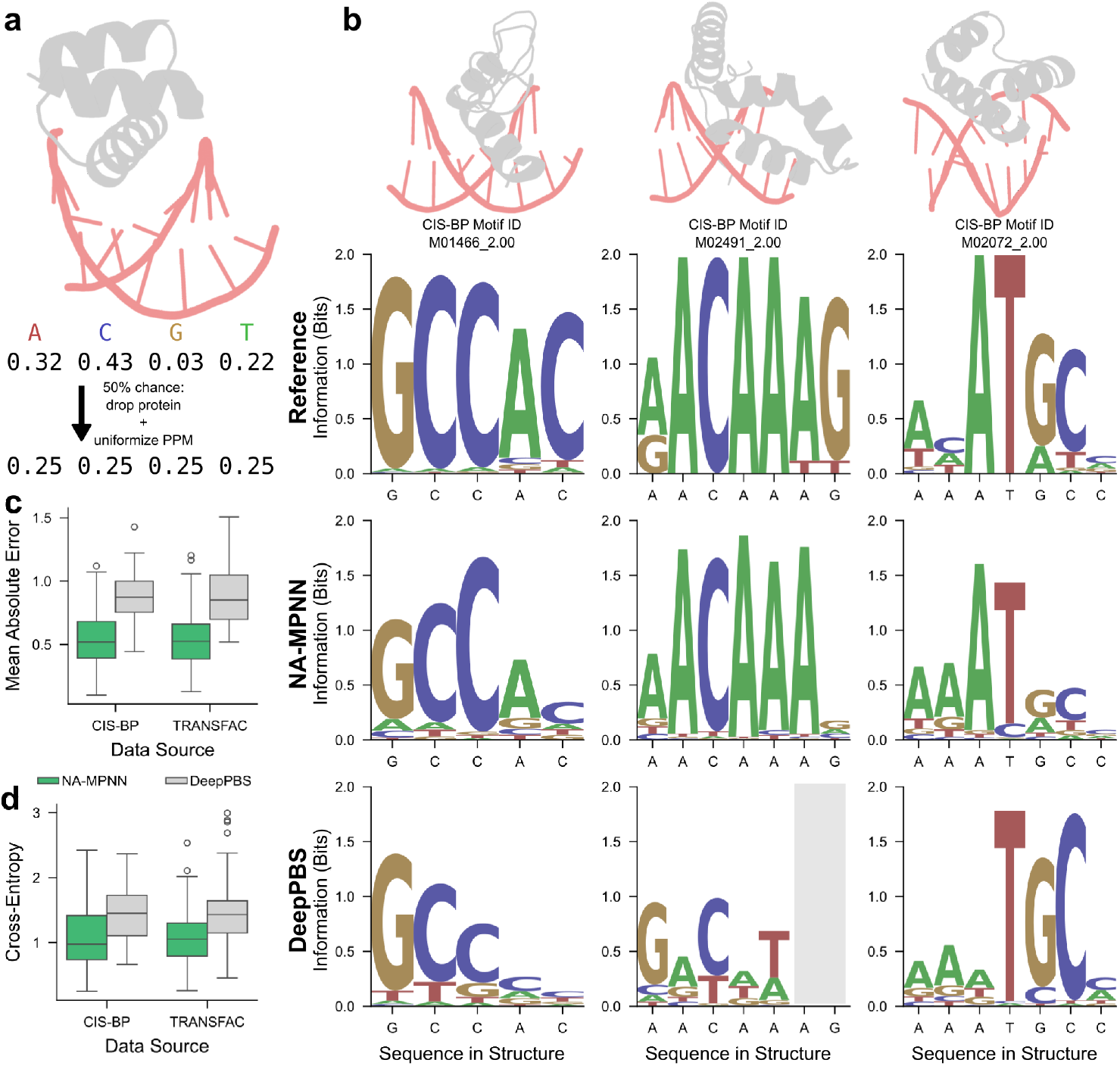
Fixed-dock protein–DNA specificity prediction. **(a)** Illustration of part of the data augmentation scheme used during training of the specificity model: randomly drop protein chains (50% probability) and substitute a uniform PPM at nucleic acid positions. **(b)** Three representative test set complexes with cartoon backbones and aligned PPMs: the experimental motif (top), NA-MPNN prediction (middle) and DeepPBS prediction (bottom), labeled with the DNA sequence from the structure. NA-MPNN more accurately reproduces the experimental base preferences. The gray box indicates positions for which DeepPBS did not make predictions. **(c–d)** Distributions of mean absolute error and cross-entropy on the CIS-BP and TRANSFAC splits of the RFNA/RFAA distillation test set: NA-MPNN attains lower medians than DeepPBS for both metrics.

## Discussion

NA-MPNN sets a new standard in the field of backbone-conditioned RNA sequence design. Given RNA structures, NA-MPNN consistently recovers 60.5% of native base identities, and secondary and tertiary structures predicted from NA-MPNN sequences using RibonanzaNet and AlphaFold 3 are closer to the input structures than those from gRNAde and RhoDesign. In the OpenKnot competition, NA-MPNN designs had the highest overall consistency measured by experimental chemical footprinting (highest median experimental OpenKnot score) among all automated methods, demonstrating the successful translation from improved *in silico* accuracy to wet-lab success. Beyond fixed-backbone sequence-design benchmarking, NA-MPNN has been coupled with a protein and nucleic acid diffusion-based backbone generator, RFDpoly^33^, to produce *de novo* RNAs and protein-DNA complexes that have been verified by electron microscopy to adopt the intended secondary and tertiary folds, underscoring the model’s utility in forward macromolecular engineering workflows.

The NA-MPNN architecture, retrained with task-specific augmentations, achieves state-of-the-art accuracy for fixed-dock protein–DNA specificity prediction. On a hold-out set of RFNA/RFAA-distilled complexes with experimental PPMs, NA-MPNN reduces the median mean absolute error and cross-entropy relative to DeepPBS despite operating solely on backbone coordinates and omitting protein side-chain atoms. This side-chain-agnostic formulation avoids information leakage about the target DNA sequence and therefore provides a stringent, early-stage filter for DNA-binding protein design^34^—well before rotamer packing or full-atom refinement. We anticipate that NA-MPNN will become a valuable component of prospective pipelines for both natural and designed transcription factors, enabling rapid, structure-aware screening of specificity profiles without the computational overhead of exhaustive docking or side-chain sampling.

In summary, NA-MPNN provides a robust and efficient tool for RNA sequence design and protein–DNA specificity prediction. We expect it to be broadly useful for creating the next generation of designed RNA molecules, transcription factors and genome engineering tools. Although not explored here, the unification of protein, DNA, and RNA inverse folding enables additional applications, including RNA-binding protein specificity prediction, backbone-conditioned sequence design of single-stranded DNA, and backbone-conditioned sequence design of protein–nucleic acid constructs.

## Methods

### RCSB Dataset

We assembled a comprehensive nucleic acid dataset from the Protein Data Bank (PDB)^22,23^, including all entries available as of January 21, 2025. Entries were retained if and only if they satisfied the following criteria: (i) heavy-atom count ≥ 100; (ii) sequence coverage ≥ 0.9; (iii) the most frequent residue was not “UNK” occurring > 20 times; (iv) resolution ≤ 3.5 Å or no resolution reported (to include NMR structures); and (v) the entry contained at least one DNA, RNA, or DNA–RNA hybrid chain. In addition, we applied an occupancy filter consistent with our model’s backbone requirements: entries were removed if no nucleic acid residue had all backbone atoms with occupancy > 0.8. Finally, we excluded any entry that failed at any point in the preprocessing pipeline (e.g., structure parsing or feature computation failures). Applying these filters yielded 15,657 preprocessed examples.

To associate experimental binding preferences with structures, we leveraged the PDB ID **+** chain ID to PPM mapping distributed with DeepPBS^21^, which links co-crystal structures to JASPAR^24^ and HOCOMOCO^25^ position probability matrices (PPMs). When a single chain mapped to multiple PPMs, we treated those PPMs as experimentally equivalent (interchangeable realizations of the same motif), and we selected among them at load time (see PPM Loading and Alignment for more details). When a single PDB had PPMs for multiple chains, we treated each chain-specific set as a distinct “unique PPM group.” This procedure produced 697 PDB-to-PPM group list mappings; a small number of referenced PPM identifiers (11) were not present in our JASPAR/HOCOMOCO snapshots and were excluded.

### RFNA/RFAA Distillation Dataset

For fixed-dock specificity prediction, we added the protein–DNA distillation complexes from RosettaFold-3^35^. This dataset includes RFNA^17^ or RFAA^18^ predictions of protein–DNA pairs drawn from CIS-BP^26^ and TRANSFAC^27^. We filtered predictions using structural-confidence thresholds of interface pAE ≤ 6 and complex pLDDT ≥ 0.85, and we removed any entry that failed downstream preprocessing. After filtering and preprocessing, the resulting dataset contained 29,460 protein–DNA complexes.

Each protein in this distillation set originates from experimental assays tied to a PPM. We paired the CIS-BP and TRANSFAC PPMs to the corresponding RFNA/RFAA complexes; when multiple PPMs were available for the same protein, we treated them as experimentally equivalent. This produced 6,267 protein–DNA distillation complexes with associated CIS-BP PPMs and 21,279 with TRANSFAC PPMs. Complexes without matched PPMs were still retained.

### Preprocessing

For every entry (and for each assembly, when applicable) across the two datasets described above, we precomputed a small set of quantities to accelerate training and standardize downstream sampling. First, we recorded per-assembly residue counts by polymer class—the total numbers of macromolecular, protein, DNA, and RNA residues—used for dynamic, token-budget-aware batching (see Training). Second, we constructed a side-chain interface mask to capture protein–nucleic acid contacts at side-chain resolution. A protein residue was marked as an interface residue if any of its side-chain atoms lay within 5 Å of a DNA/RNA side-chain atom, and a nucleic acid residue was marked as an interface residue if any of its side-chain atoms lay within 5 Å of a protein side-chain atom. For efficiency, distance checks were limited to the 48 nearest neighbors of each residue, using Cα as the representative atom for proteins and C1′ for nucleic acids. These masks are later used for training-time augmentations (see Data Augmentation Details).

### Clustering

We clustered the sequence of every protein chain in both datasets using CD-HIT^36–38^ with parameters: -c 0.4 -n 2 -d 0 -M 16000 -T 0 -aL 0.9 -aS 0.9. This produced 6,658 clusters with 83 sequences remaining unclustered. For nucleic acid chains, we applied CD-HIT-EST^39^ after standardizing sequences to a shared DNA/RNA representation: U mapped to T and all non-canonical residues mapped to X (for clustering only). We used parameters: -c 0.8 -n 4 -d 0 -M 16000 -T 0 -l 4 -aL 0.9 -aS 0.9, yielding 9,849 clusters with 176 sequences unclustered.

Cluster assignments serve two roles downstream: (i) defining train/validation/test holdouts (see Train/Validation/Test Split), and (ii) controlling sampling probability during training via rarity-aware weights (see Training). Entries for which all protein and all nucleic acid chains lacked cluster assignments were removed; this step eliminated 24 structures from the RCSB dataset.

### Train/Validation/Test Split

We constructed two datasets aligned to the two learning objectives: (i) a design dataset for backbone-conditioned sequence design and (ii) a specificity dataset for fixed-dock protein–DNA base-preference prediction. The design dataset was derived exclusively from the RCSB crystal structures described above and contains 15,633 entries spanning 6,137 protein chain clusters and 6,092 nucleic acid chain clusters. Because this model is used primarily for nucleic acid design, we split by nucleic acid chain clusters. We additionally held out all nucleic acid chain clusters corresponding to a pseudoknot test set for *in silico* evaluation, defined by the PDB IDs: 1DRZ, 2M8K, 2MIY, 3Q3Z, 4OQU, 4PLX, 4ZNP, 7KD1, 7KGA, 7QR4 (borrowed from the OpenKnot challenge^31^). From the remaining nucleic acid clusters we randomly assigned 10% to test and 10% to validation, after excluding clusters with degree > 25 to prevent a few large families from dominating the held-out sets. This yielded 4,864 training clusters, 609 validation clusters, and 619 test clusters. Because examples are full complexes while clustering is at the chain level, we enforced a leak-free assignment rule: an example is placed in test if any of its nucleic acid chains maps to a test cluster; otherwise in validation if any maps to a validation cluster; otherwise in train. The resulting split is summarized in Supplementary Table S1.

The specificity dataset combines the RCSB crystal structures with the RFNA/RFAA distillation set, for a total of 45,093 entries encompassing 6,658 protein chain clusters and 9,849 nucleic acid chain clusters. Using the same degree cap (≤ 25) and 10%/10% validation/test proportions, we split by protein chain clusters (reflecting the protein-centered notion of specificity). This produced 5,328 training clusters, 665 validation clusters, and 665 test clusters. Example-level assignment followed the analogous leak-prevention rule: test if any protein chain lands in a test cluster; else validation if any lands in a validation cluster; else train. The resulting split is summarized in Supplementary Table S2.

### PPM Loading and Alignment

When training the specificity model, we load experimental position probability matrices (PPMs) whenever they are available for a given complex. If an example is associated with multiple experimental PPMs that belong to the same unique PPM group (i.e., experimentally equivalent motifs drawn from different sources), we randomly sample one of the experimentally equivalent PPMs each time the example is loaded, so that training stochastically explores assay variability across epochs. Because a complex can contain multiple unique PPM groups (e.g., multiple interfaces), multiple PPMs may be present for the same structure during a given load (we load one PPM per “unique PPM group”). For every sampled PPM, we compute its reverse complement and align both the original and reverse-complement PPMs to every nucleic acid chain in the complex using a modified version of the DeepPBS^21^ PPM–structure alignment procedure. Specifically, for each chain we enumerate all possible overlaps of length ≥ 5 and compute the sum of per-position information-content-weighted Pearson correlation coefficient (IC-weighted PCC) at each offset. Mathematically, per-position information content is calculated as:

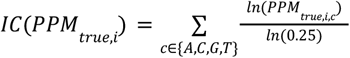

where *PPM*_*true,i,c*_is the experimental probability for the *i* ^*th*^ position and residue type *c*, after adding an epsilon of 10^−10^ to all residue type probabilities for the *i*^*th*^ residue and renormalization. Utilizing this formula in the IC-weighted PCC computation:

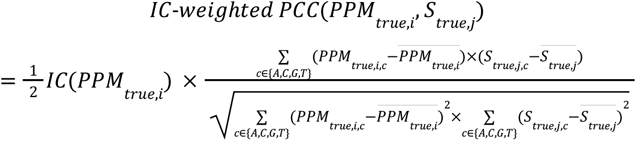

where *PPM*_*true,i*_ is the PPM column for the *i* ^*th*^ position, *S*_*true,j*_ is the one-hot true sequence for the *j*^*th*^ residue, and ⨪ indicates the mean across the residue type index *c*. When the PPM column is uniform, the denominator vanishes; in these cases, we define

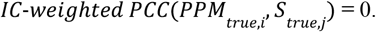

Non-canonical DNA residues in the structure are loaded as the unknown nucleic acid token; positions that are unknown are excluded from both the IC-weighted PCC calculation and the minimum overlap-length check. For each PPM we record the best-scoring alignment (highest per-position IC-weighted PCC sum) together with its chain and offset; ties are retained as multiple best alignments (useful, for example, when the same motif appears twice in one complex). We then write the aligned PPM columns for the best alignment(s) into a global, per-complex aligned PPM representation and repeat until all PPMs have been incorporated. If two aligned PPMs overlap on the same sequence position, we resolve the conflict by retaining the column with the higher local alignment score; if the underlying sequence position is unknown, we break ties by selecting the column with the higher information content.

### Additional Data Featurization and Architecture Details

We follow the ProteinMPNN convention of retaining a residue only if all backbone atoms have occupancy > 0.8. In addition to the architectural details discussed in Results: NA-MPNN Architecture, we provide the following details on node embeddings and pairwise distance calculations. The polymer-type node features are constructed as follows: for each residue we encode a one-hot of polymer type (protein, DNA, RNA, unknown), project it with a bias-free linear layer to the model hidden dimension (matched to edge features), apply LayerNorm, and map with a second linear layer (with bias) to obtain the final node embedding. We represent each residue with coordinates for the combined set of protein and nucleic acid backbone atoms; only the subset corresponding to the residue’s polymer type is populated. During construction of residue–residue distance features, we compute pairwise distances over this unified set but zero out the radial-basis embeddings for atom pairs that do not exist. This yields a simple, unified representation for proteins, DNA, and RNA while avoiding leakage of non-physical information.

### Data Augmentation Details

We employed three augmentations only when training the specificity model to emphasize protein-induced preferences and to incorporate examples lacking experimental motifs. (i) For nucleic acid–only complexes (DNA, RNA, or DNA–RNA hybrids), we treated every nucleic acid position as carrying a uniform PPM (0.25, 0.25, 0.25, 0.25) over the four canonical bases. (ii) For protein–nucleic acid complexes, we drop all protein chains with probability 0.5 and replace the PPM at every nucleic acid position with the uniform PPM, encouraging the model to attribute specificity to protein context rather than nucleic acid geometry alone. (iii) Using the side-chain interface mask described above, we uniformized any non-interface nucleic acid residue that lacked an experimental PPM; residues at the side-chain interface without experimental PPMs retained a label-smoothed one-hot PPM derived from the crystal sequence. Together, these procedures provide information-free supervision away from the interface while preserving motif signal at the interface.

### Loss

Optimization follows ProteinMPNN^28^ with two task-specific modifications. First, we apply label smoothing with coefficient ϵ = 0.1 within polymer classes only: probability mass is redistributed over protein tokens for proteins and over nucleic acid tokens for DNA/RNA, preventing leakage across polymer types during training. Second, during specificity training, whenever an experimental PPM is available at a position, we swap the ground-truth one-hot with that PPM column in the cross-entropy target, thus training the model to match distributions rather than single bases where assays provide positional preferences. The loss can be summarized as:

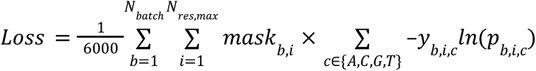

where *N*_*batch*_ is the number of batches, *N*_*res,max*_ is the maximum number of residues across the batch, *mask*_*b,i*_ is 1 if the residue in example *b* at residue index *i* exists and was designed, *y*_*b,i,c*_ is the label-smoothed true label (one-hot sequence or experimental PPM) for example *b* at residue index *i* for residue type *c, ln*(*p*_*b,i,c*_) is the model’s log probability for example *b* at residue index *i* for residue type *c*.

### Training

The per-example sampling weight is computed as follows:

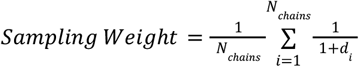

where *N*_*chains*_ is the number of protein and nucleic acid chains, and *d*_*i*_ is the degree of the corresponding chain’s sequence cluster after subsetting to the design or specificity dataset (see Train/Validation/Test Split). This sampling procedure upweights examples containing protein or nucleic acid chains from rare sequence clusters.

As in LigandMPNN^29^, we train with the Adam optimizer (β_1_ = 0.9, β_2_ = 0.98, ϵ = 10^-9^) using token-budgeted batches of 6,000 tokens, automatic mixed precision, gradient-norm clipping of 1.0, and gradient checkpointing on a single NVIDIA A100 GPU. Both the design and specificity models are trained for 100,000 batches (training curves in Supplementary Figure S1); the final checkpoints used in evaluation were selected earlier than the maximum step based on validation behavior.

### Evaluation Datasets

For both design and specificity tasks, we applied additional filtering to the validation and test partitions to standardize inputs across tools and ensure leak-free evaluation. For the design sets, we retained a complex only if all of its nucleic acid chains belonged to the corresponding validation or test nucleic acid chain clusters defined by the split (see Train/Validation/Test Split). For the specificity sets, we enforced the analogous constraint on protein chains (all protein chains must fall in the corresponding validation or test protein clusters). We then removed any example with less than 20 or greater than 1000 total polymer residues (counting protein, DNA, and RNA) and excluded cases where CIF to PDB conversion failed (several downstream tools accept PDB only), yielding the finalized evaluation datasets used throughout. We further required model applicability: an example was only kept if all relevant models successfully executed on the example.

The design test set was further partitioned into two specialized benchmarks: 1) the RNA-monomer test set (complexes containing exactly one RNA chain and no additional chains), and 2) the pseudoknot test set (a set of 10 pseudoknot PDB ID [chain ID] pairs discussed in Train/Validation/Test Split above: 1DRZ [B], 2M8K [A], 2MIY [A], 3Q3Z [A], 4OQU [A], 4PLX [A], 4ZNP [A], 7KD1 [A], 7KGA [A], 7QR4 [B]). To standardize inputs for these subsets, we sourced structures from RNAsolo^40^ rather than directly from the PDB. RNAsolo only contains RNA chains, which has the effect of removing protein chains in some pseudoknot cases. When multiple RNAsolo sources existed, we preferred the Rfam version, defaulting to BGSU if no Rfam structure was available—except for PDB IDs 1VC5 and 4ZNP, where we prioritized BGSU due to downstream processing issues with the Rfam files. The resulting design validation and test sets used for benchmarking are summarized in Supplementary Table S3.

For specificity, we retained only complexes with matched experimental PPMs. We additionally split validation and test entries by structure and experimental PPM source, distinguishing crystal structure, RFNA/RFAA distillation structure with CIS-BP PPM, and RFNA/RFAA distillation structure with TRANSFAC PPM. The final specificity validation and test sets are reported in Supplementary Table S4.

### Evaluation Metrics

For sequence design, the primary metric is sequence recovery—the fraction of designed positions whose identities match the native residue in the crystal structure:

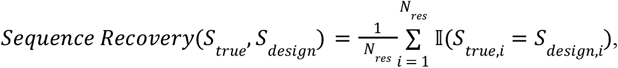

where *N*_*res*_ is the total number of residues, *S*_*true,i*_ and *S*_*design*.*i*_ are the true and designed residue type at residue index *i*, and the indicator function II (*S*_*true,i*_ = *S*_*design*.*i*_) is 1 if the true and designed residue types are equal and 0 otherwise. To avoid discrepancies arising from residues that any given tool failed to load or design, we use each tool’s own reported sequence-recovery procedure, and we restrict computation to designed residues only. Sequence recovery is computed for the design validation set, the design test set, and the RNA-monomer and pseudoknot subsets.

On the pseudoknot subset, we evaluate additional structure- and reactivity-based metrics. DSSR v2.3.2^41^ is used to extract the ground-truth secondary structure from the native crystal structures. For each designed sequence, RibonanzaNet predicts 2A3 reactivity profiles, from which we compute predicted OpenKnot scores (see https://github.com/eternagame/OpenKnotScore)^31^ using the predicted reactivity together with the DSSR ground truth. AlphaFold 3^19^ is run with one seed, five diffusion samples, and no MSAs or templates; we record the mean pLDDT over all atoms. Predicted models are superimposed with Biotite^42^ onto the native structure to compute C1′-RMSD.

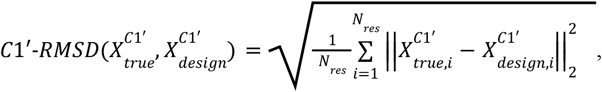

where *N*_*res*_ is the total number of residues, 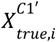 is the C1′ coordinate for the *i*^*th*^ residue from the true structure,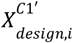 is the C1′ coordinate for the *i*^*th*^ residue of the designed sequence from the AlphaFold 3-predicted structure (following superimposition to the true structure), and ∥·∥_2_ represents the vector length. When designed and native sequences differ in length (e.g., terminal residues not designed by a method), we enumerate all valid overlaps, select the best by C1′ RMSD, and crop sequence, coordinates, and secondary structure accordingly (if one base of a native pair is cropped, its remaining partner is set to a loop). Additional secondary structure, tertiary structure, and structure confidence metrics are reported in Supplementary Fig. S4.

For fixed-dock protein–DNA specificity, for each tool, we load the first experimentally equivalent PPM for every unique PPM group associated with a complex (i.e., no randomization at evaluation). We generate the reverse-complement PPM and align both the original and reverse complement to the tool’s loaded DNA chain sequences using the same IC-weighted PCC alignment procedure as in training (see PPM Loading and Alignment). We then subset the aligned experimental and predicted PPMs to positions where both exist. At each retained position, we compute mean absolute error (MAE) and cross-entropy over the four DNA bases, and finally average along the aligned window to obtain per-example scores. Represented mathematically:

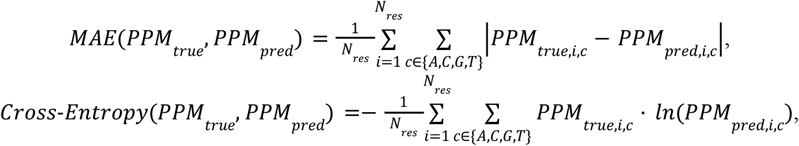

where *N*_*res*_ is the total number of residues, *PPM*_*true,i,c*_and *PPM*_*pred*.*i,c*_ are the true and predicted probability for the *i*^*th*^ residue and residue type *c*.

### Model and Hyperparameter Selection

For the design model, we selected the evaluation checkpoint using the design validation set. For a panel of checkpoints (Supplementary Fig. S2), we computed sequence recovery separately for four contexts—DNA-only, RNA-only, DNA with fixed protein context, and RNA with fixed protein context—for every example in the validation set and reported the median per context. The checkpoint after 19,137 training batches achieved the best aggregate balance across these four classes and was chosen for all subsequent design evaluations.

For the specificity model, we used the distillation validation set to select both the checkpoint and inference hyperparameters. For each candidate checkpoint we evaluated MAE and cross-entropy on all distillation validation examples and reported the median of each metric (Supplementary Fig. S2). The checkpoint after 70,114 training batches provided the optimal trade-off and was selected. We then performed a grid search over the sampling temperature and number of stochastic samples drawn from the model at inference; 30 samples and temperature = 0.6 yielded balanced performance on MAE and cross-entropy (Supplementary Fig. S3) and were adopted for testing.

### Model Evaluation

For the design task, we generated 10 sequences per backbone at temperature = 0.1 and reported metrics over all designed sequences. For NA-MPNN, residues missing any backbone atom were not designed; when protein chains were present (validation and overall test sets), the protein was held fixed and served as context. gRNAde^15^ was run with the “Das” data split and one conformer, and RhoDesign^16^ was run without secondary structure conditioning to enable a tertiary structure inverse-folding comparison. Only NA-MPNN was executed on the full validation and whole test sets; all three methods (NA-MPNN, gRNAde, RhoDesign) were executed on the RNA-monomer and pseudoknot subsets.

For the specificity task, on the validation set, we evaluated NA-MPNN only. For testing, NA-MPNN and DeepPBS^21^ were run on the RFNA/RFAA distillation test set, using the alignment and scoring protocol described under Evaluation Metrics. Analyses for the crystal structure subset are provided in Supplementary Fig. S6.

### Design and Experimental Validation of RNA Pseudoknots

For Round 6 of the OpenKnot challenge^31^, we generated NA-MPNN sequences for the 12 fixed-backbone pseudoknot puzzles. We ranked candidates by RibonanzaNet-predicted secondary structure consistency, computing an F_1_ score between predicted and target secondary structures (see Supplementary Information: Additional *In Silico* Metrics for RNA Pseudoknot Sequence Design). The OpenKnot challenge organizers experimentally evaluated the submitted sequences and computed experimental OpenKnot scores from the SHAPE-derived reactivity profiles relative to the target secondary structure. Experimental results were plotted for designs with high quality experimental reactivity profiles (SN_filter = 1)^31^. NA-MPNN performance on Rounds 5, 7a, and 7b of the challenge is summarized in Supplementary Fig. S5.

### Software Dependencies

All modeling and analysis was implemented in Python, using NumPy^43^ and pandas^44,45^ for data preprocessing; NumPy, ProDy^46–48^, and PyTorch^49^ for featurization and data loading; and PyTorch for model training. Biotite^42^ supported structural superposition and computation of RMSD/LDDT/GDDT. NumPy, pandas, matplotlib^50^, seaborn^51^, Logomaker^52^, and PYMOL^53^ were used for data analysis and figure generation.

## Code Availability

Training and inference code for NA-MPNN—along with data-preprocessing, evaluation, and visualization notebooks; model weights; installation instructions; and usage demonstrations—is available on GitHub: https://github.com/baker-laboratory/NA-MPNN.

## Data Availability

For the design model, the PDB IDs of all structures used for training and evaluation are provided in the repository: https://github.com/baker-laboratory/NA-MPNN.

For the specificity model, source/ID pairs for non-TRANSFAC PPMs used in training and evaluation are listed at the same repository. Associated PDB IDs for relevant structures are included, and test-set RFNA/RFAA distillation structures derived from CIS-BP are provided. TRANSFAC-related data cannot be redistributed due to licensing restrictions.

Specifications for the OpenKnot challenge puzzles from Rounds 5, 6, 7a, and 7b can be found at https://eternagame.org/challenges/11843006. Experimental data for OpenKnot designs are available at https://github.com/eternagame/OpenKnotAIDesignData; rounds 1, 2, 3, and 4 in this dataset CSV correspond to Eterna OpenKnot rounds 5, 6, 7a, and 7b respectively.

## Author Contributions

Ideation: A.K., A.F., L.M., R.M., R.P., J.D., C.G., D.B.; Development of NA-MPNN: A.K., J.D.; RFNA/RFAA distillation set generation: L.M.; RNA sequence design and evaluation for OpenKnot: A.K., A.F.; *In silico* evaluation of NA-MPNN: A.K., C.G.; Figure creation: A.K., R.M., A.F.; Writing: A.K., D.B., R.M.; Editing: A.K., A.F., L.M., R.M., R.P., J.D., C.G., D.B.; Supervision: D.B.

## Acknowledgements

We thank the Eterna project and the Das lab (Stanford, HHMI) for coordination and experimental evaluation of the Eterna OpenKnot challenge; Frank DiMaio, Phil Bradley, Brian Trippe, Paul Kim, Han Raut Altae-Tran, Pascal Sturmfels, Enisha Sehgal, and Yuliya Politanska for helpful methodological discussions; Luki Goldschmidt for maintaining the computational resources utilized for this research; and Madison Kennedy for editorial assistance.

This work was supported by The Audacious Project (C.G.); The Open Philanthropy Project (J.D.); the Gates Foundation, grant no. INV-043758 (A.F., R.M.); the National Institute for Allergy and Infectious Diseases, NIH grant no. 1U19AI181881-01 (A.K.); the Washington Research Foundation (C.G.); the Defense Threat Reduction Agency, grant no. HDTRA1-22-1-0012 (L.M.); Howard Hughes Medical Institute (D.B.). This work was based upon work supported by the National Science Foundation, grant no. CHE-2226466 (L.M., R.P., C.G., D.B.).

## Notes

### Competing Interest Statement

The authors have declared no competing interest.

### Summary of Updates

Minor grammatical edits; revision of Methods: RFNA/RFAA Distillation Dataset section.

