## Supplementary Information for "RNA sequence design and protein–DNA specificity prediction with NA-MPNN"

*Additional In Silico Metrics for RNA Pseudoknot Sequence Design*

Using DSSR[1] to extract the ground-truth secondary structure from native crystal structures and RibonanzaNet[2] to predict secondary structure for each designed sequence, we computed $F_1$ scores for both base pairs and loops. For pair $F_1$, we computed the following:

$$Pair\ F_1 = \frac{2 \times TP_{pairs}}{2 \times TP_{pairs} + FP_{pairs} + FN_{pairs}},$$

where true positives ($TP_{pairs}$) are residue–residue base pairs present in both native structures and design predictions; false positives ($FP_{pairs}$) are base pairs present only in the design prediction; and false negatives ($FN_{pairs}$) are native base pairs missing from the design prediction. For loop F1:

$$Loop\ F_1 = \frac{2 \times TP_{loops}}{2 \times TP_{loops} + FP_{loops} + FN_{loops}},$$

where true positives ($TP_{loops}$) are loop positions found in both native structures and design predictions, whereas false positives ($FP_{loops}$) and false negatives ($FN_{loops}$) are loop positions unique to the design prediction or to the native structure, respectively. In addition to the AlphaFold 3[3] metrics described in Methods: Evaluation Metrics, we also recorded mean pAE (averaged over residue pairs) and pTM. Biotite[4] was used to compute C1′-LDDT and C1′-GDDT from the AlphaFold 3-predicted design models. As summarized in Supplementary Fig. S4, NA-MPNN exhibits superior secondary and tertiary structure recovery and predicted confidence across these additional metrics.

*Additional Wet-Lab Performance for RNA Pseudoknot Sequence Design*

NA-MPNN sequences were generated for the puzzles from Rounds 5, 7a, and 7b, as described in Methods: Design and Experimental Validation of RNA Pseudoknots. The rounds differed primarily in the filtering pipeline. In Round 5, RibonanzaNet was unavailable, so EternaFold[5]-predicted secondary structures were used instead. During Rounds 7a and 7b, we augmented filtering with RibonanzaNet-predicted OpenKnot scores, as well as Chai-1[6]/AlphaFold 3-predicted secondary ($F_1$ score) and tertiary structure (TM-score[7], C1′-GDDT) consistency.

The experimental results in Figure 3 correspond to Round 6 of the OpenKnot competition[2]. In Round 5, NA-MPNN demonstrated a similar trend; NA-MPNN sequences achieved equivalent or higher OpenKnot scores compared to starting sequences and other sequence sources. In later rounds, NA-MPNN lagged behind the starting (wild-type) sequences and player-designed Eterna sequences; a plausible contributing factor is that several targets were computationally generated backbones whose quality was lower than that of crystal structure backbones used in earlier rounds. These results are displayed in Supplementary Fig. S5.

*Additional Fixed-Dock Protein–DNA Specificity Evaluation*

Supplementary Fig. S6 reports results on the crystal structure test subset. Fig. S6c–d shows that NA-MPNN underperforms DeepPBS on this subset. The primary explanation for this discrepancy is split strategy: DeepPBS[8] uses a random train/test split that samples (without replacement) from the same protein-sequence clusters, whereas our protocol holds out entire protein sequence clusters (40% sequence identity) for testing (see Train/Validation/Test Split). Consequently, the crystal structures in the test set comprise proteins entirely unseen by NA-MPNN during training but potentially within-cluster for DeepPBS, accounting for the performance gap. Notably, Fig. S6a–b highlight crystal cases where NA-MPNN outperforms DeepPBS: (i) a complex presenting two instances of the same binding motif, and (ii) a complex with a non-standard DNA helix where DeepPBS fails to detect the helix and thus cannot produce predictions at those positions. Because NA-MPNN does not rely on single-helix detection, it can generate predictions for irregular or single-stranded DNA regions. By construction—via shared nucleic acid tokens—the same machinery could, in principle, be extended to protein–RNA specificity predictions.

| Nucleic Acid Type | Protein? | Train | Validation | Test |
|---|---|---|---|---|
| DNA | ✘ | 2015 | 151 | 147 |
| | ✔ | 5660 | 483 | 527 |
| RNA | ✘ | 1451 | 169 | 160 |
| | ✔ | 2796 | 397 | 394 |
| DNA/RNA Hybrid | ✘ | 59 | 0 | 4 |
| | ✔ | 43 | 2 | 0 |
| Multi-NA Types | ✘ | 92 | 3 | 12 |
| | ✔ | 814 | 125 | 129 |

**Table S1. Design datasets overview.**
The example counts for the design train/validation/test datasets, broken down into examples containing only DNA, only RNA, only DNA/RNA hybrid chains, or multiple nucleic acid types with or without protein.

| Structure Source | Experimental PPM? | Train | Validation | Test |
|---|---|---|---|---|
| Distillation (CIS-BP) | ✘ | 829 | 5 | 2 |
| | ✔ | 5982 | 163 | 122 |
| Distillation (TRANSFAC) | ✘ | 1066 | 9 | 3 |
| | ✔ | 21029 | 137 | 113 |
| Crystal | ✘ | 12731 | 903 | 1385 |
| | ✔ | 485 | 46 | 83 |

**Table S2. Specificity datasets overview.**
The example counts for the specificity train/validation/test datasets, categorized by structure source and availability of experimental PPMs.

| Nucleic Acid Type | Protein? | Validation | Test | RNA-Monomer Test | Pseudoknot Test |
|---|---|---|---|---|---|
| DNA | ✘ | 41 | 39 | 0 | 0 |
| | ✔ | 229 | 237 | 0 | 0 |
| RNA | ✘ | 105 | 88 | 63 | 10 |
| | ✔ | 131 | 139 | 0 | 0 |

**Table S3. Evaluation datasets for design models.**
The final example counts in the evaluation datasets for the nucleic acid sequence-design models, split by nucleic acid type and the presence of protein. Note, two of the RNA pseudoknot examples contained protein chains, which were removed in the creation of the pseudoknot test set.

| Structure Source | Validation | Test |
|---|---|---|
| Distillation (CIS-BP) | 163 | 118 |
| Distillation (TRANSFAC) | 137 | 110 |
| Crystal | 18 | 46 |

**Table S4. Evaluation datasets for specificity models.**
The final example counts in the evaluation datasets for the fixed-dock protein–DNA specificity prediction models, split by structure source.

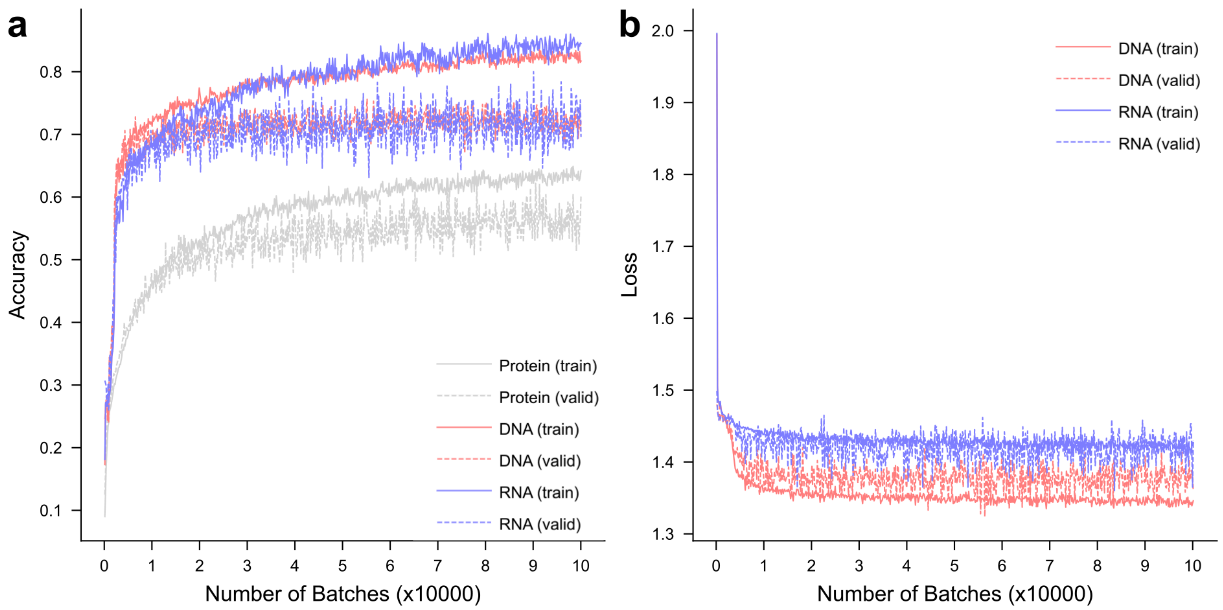


**Figure S1. Training curves for NA-MPNN design and specificity models.**
**(a)** The training curve for the NA-MPNN design model. Accuracy (sequence recovery) is plotted against batch number for protein, DNA, and RNA residues on the training and held-out validation sets **(b)** The training curve for the NA-MPNN specificity model. Loss is plotted against the batch number for DNA and RNA residues on the training and held-out validation sets; loss for the specificity model accounts for experimental PPMs when available.

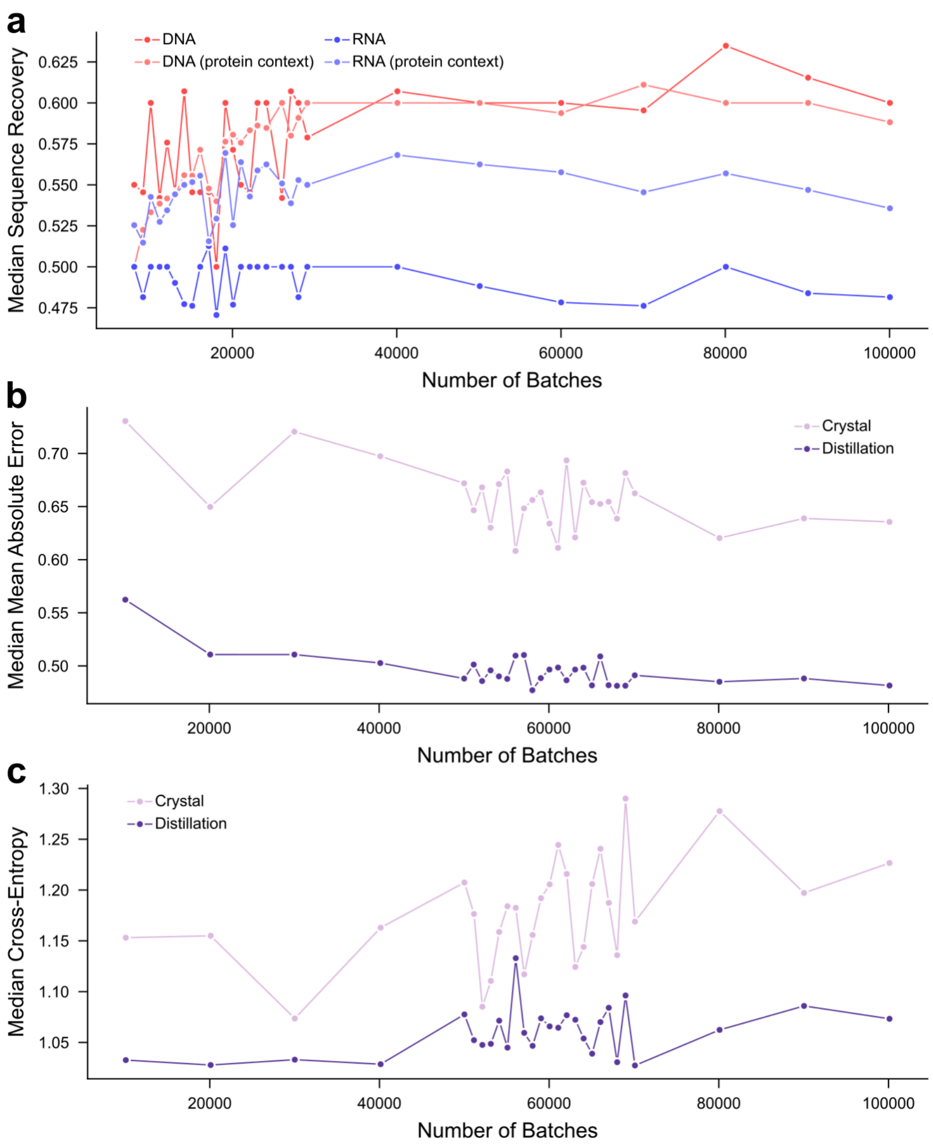


**Figure S2. Checkpoint sweep for NA-MPNN design and specificity models.**
**(a)** Median sequence recovery of DNA-only, RNA-only, DNA in fixed protein context, and RNA in fixed protein context, computed on the design validation set for various checkpoints of the NA-MPNN design model. The checkpoint after 19,137 batches provided the best tradeoff between evaluation contexts. **(b–c)** Median MAE and median cross-entropy between experimental and predicted PPMs, computed on the specificity validation set for various checkpoints of the NA-MPNN specificity model. The checkpoint after 70,114 batches achieved the best balance of the two metrics.

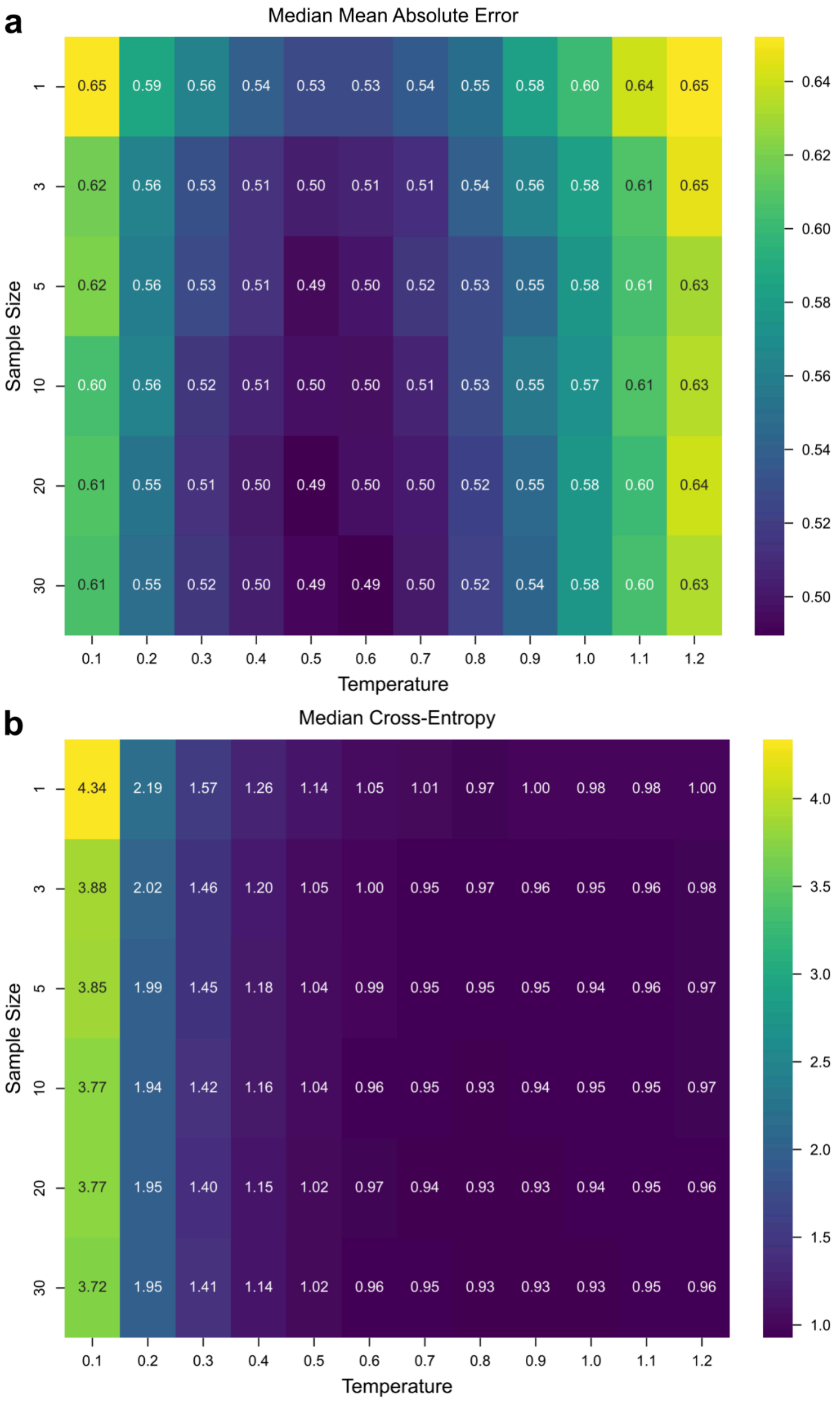


**Figure S3. Inference hyperparameter grid search for NA-MPNN specificity model.**
**(a–b)** MAE and cross-entropy grid searches over number of samples and sampling temperature, computed on the specificity distillation validation set for the NA-MPNN specificity model. A sample size of 30 and sampling temperature of 0.6 were selected for model evaluation.

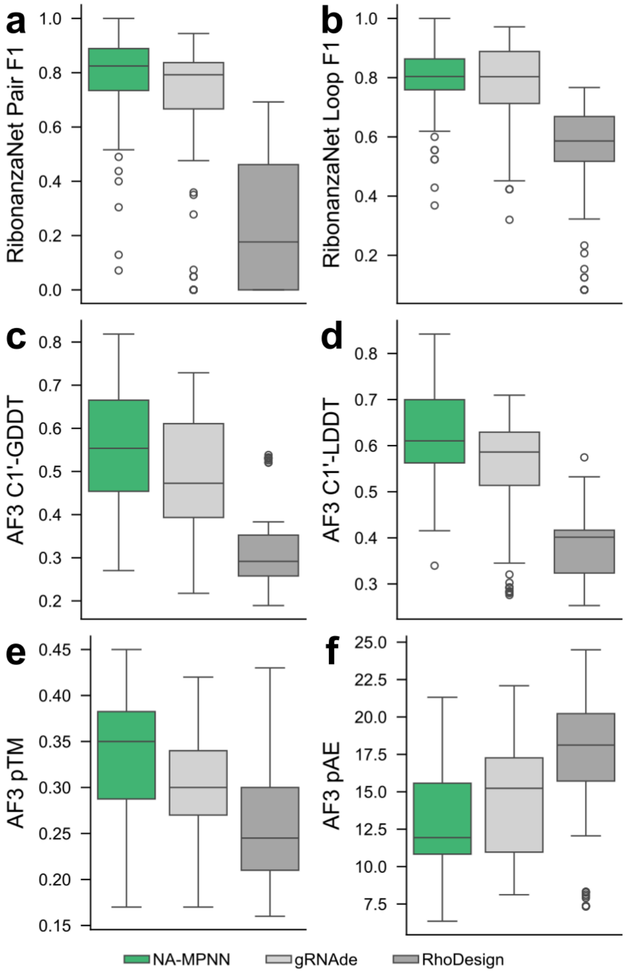


**Figure S4. Additional *in silico* metrics for RNA pseudoknot test set.**
**(a–b)** RibonanzaNet-predicted secondary structure pair and loop $F_1$ score distributions on the RNA pseudoknot test set; NA-MPNN achieves the best secondary structure recovery. **(c–d)** C1′-GDDT and C1′-LDDT distributions computed from AlphaFold 3 predictions of designed pseudoknot sequences; the predicted tertiary structure of NA-MPNN-designed sequences matches the native backbone more closely than competing methods. **(e–f)** AlphaFold 3 pTM and pAE distributions across designed RNA pseudoknot sequences; NA-MPNN sequences result in more confident AlphaFold 3 predictions.

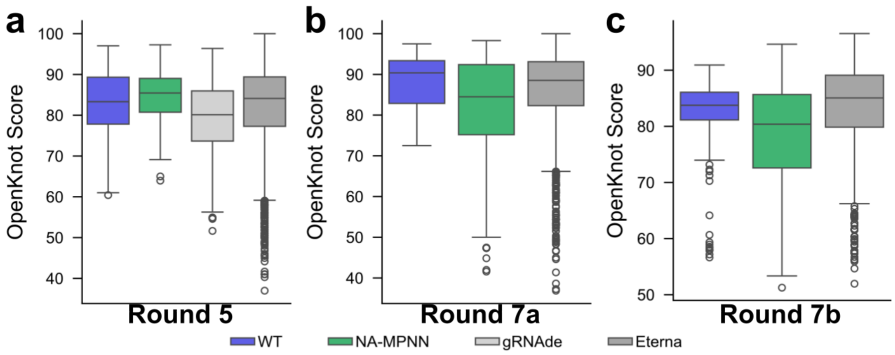


**Figure S5. Additional experimental evaluation of NA-MPNN pseudoknot sequences.** **(a–c)** Distribution of experimentally determined OpenKnot scores across all puzzles in OpenKnot Rounds 5, 7a, and 7b for the starting sequences (WT), NA-MPNN sequences, gRNAde sequences (when available), and Eterna player-created sequences. In Round 5, NA-MPNN achieves the highest median score. Several puzzles in Rounds 7a and 7b consisted of lower quality, computationally generated pseudoknot backbones (as opposed to the crystal structures from Rounds 5 and 6); due to the dependence of NA-MPNN on high quality backbones, median performance lags behind starting sequences and the Eterna player-created sequences.

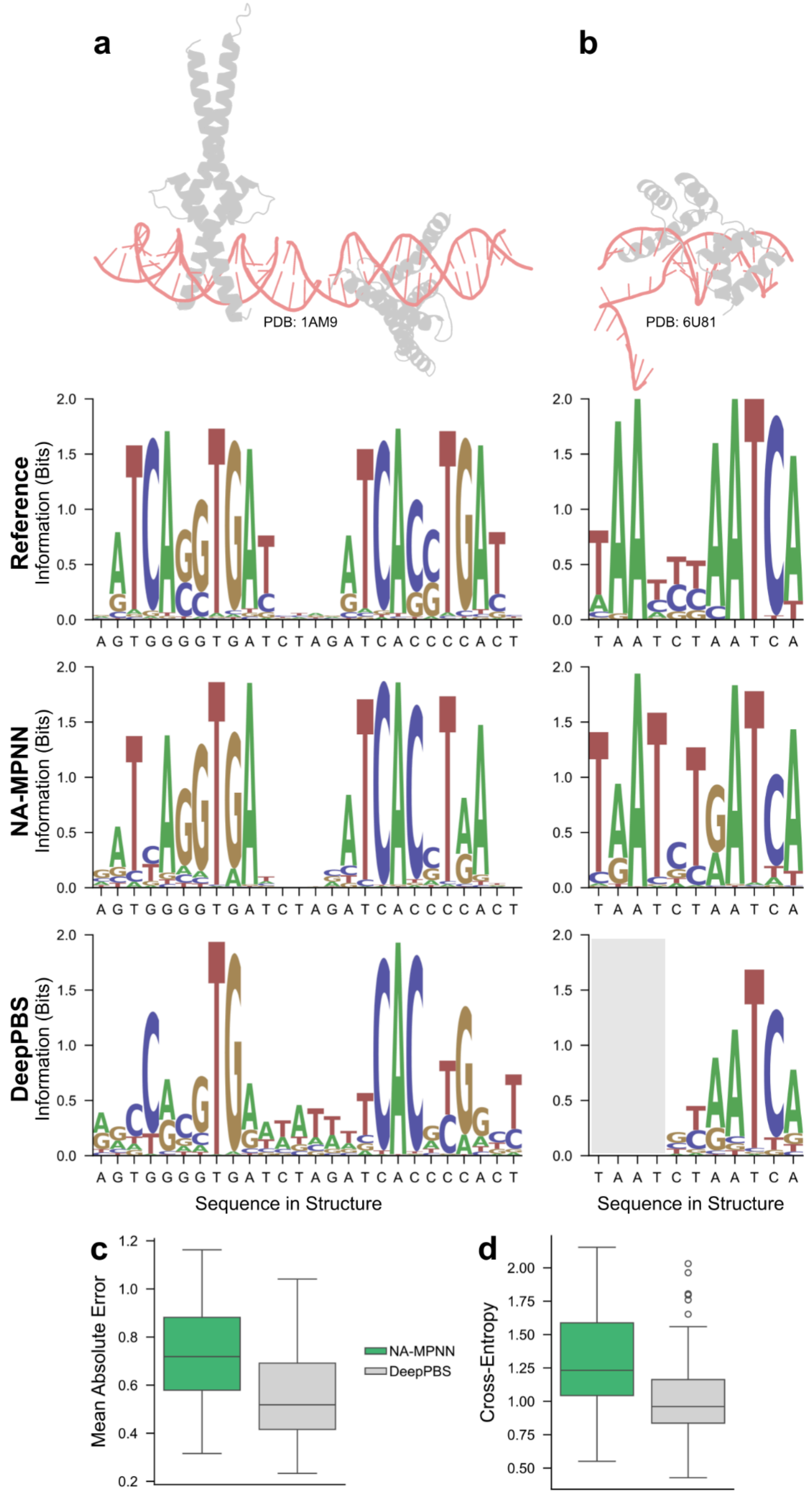
a
b
PDB: 1AM9
PDB: 6U81
Reference
NA-MPNN
DeepPBS
Information (Bits)
Sequence in Structure
c
d
Mean Absolute Error
Cross-Entropy
NA-MPNN
DeepPBS

**Figure S6. Fixed-dock protein–DNA specificity prediction on crystal structure test set.** **(a–b)** Two representative examples from the crystal structure split of the test set with cartoon backbones and aligned PPMs: the experimental motif (top), NA-MPNN prediction (middle), and DeepPBS prediction (bottom), labeled with the DNA sequence from the structure. The gray box indicates positions for which DeepPBS did not make predictions. **(c–d)** Distribution of MAE and cross-entropy on the crystal structure test set split.
